# Dissecting the roles of MBD2 isoforms in regulating NuRD complex function during cellular differentiation

**DOI:** 10.1101/2021.03.17.435677

**Authors:** Nina Schmolka, Jahnavi Bhaskaran, Ino D. Karemaker, Tuncay Baubec

## Abstract

The Nucleosome Remodelling and Deacetylation (NuRD) complex is a crucial regulator of cellular differentiation. Two members of the Methyl-CpG-binding domain (MBD) protein family, MBD2 and MBD3, are known to be integral, but mutually exclusive subunits of the NuRD complex. Several MBD2 and MBD3 isoforms are present in mammalian cells, resulting in distinct MBD-NuRD complexes. If these different complexes serve distinct biochemical and/or functional activities during differentiation is not completely understood. Based on the essential role of MBD3 in lineage commitment, we systematically investigated a diverse set of MBD3 and MBD2 variants for their potential to rescue the differentiation block observed in mouse embryonic stem cells (ESCs) lacking MBD3. Our study reveals that while MBD3 is indeed crucial for ESC differentiation to neuronal cells, this function is independent of its MBD domain or binding to methylated DNA. While MBD3 isoforms are highly redundant, we identify that MBD2 isoforms vary in their potential to fully rescue the absence of MBD3 during lineage commitment. Full-length MBD2a only partially rescues the differentiation block; MBD2b, which lacks the N-terminal GR-rich repeat, fully rescues the differentiation block in MBD3 KO ES cells, and cells expressing the testis-specific isoform MBD2t that lacks the coiled-coil domain required for NuRD interactions are not able to generate any differentiated cells. In case of MBD2a, we further show that removing the m-CpG DNA binding capacity or the GR-rich repeat renders the protein fully redundant to MBD3, highlighting the requirements for these domains in diversifying NuRD complex function. In sum, our results highlight a partial redundancy of MBD2 and MBD3 during cellular differentiation and point to specific functions of distinct MBD2 isoforms and specific domains within the NuRD complex.

## Introduction

Cellular differentiation entails establishment of new cell identities through changes in transcriptional programs. Multiple components involving external stimuli, transcription factors and chromatin modifications play important roles in orchestrating gene expression during developmental transitions. Chromatin remodelling complexes are key components of this process, enabling the change of chromatin structure and the accessibility of specific genomic sites *(1)*. The Nucleosome Remodelling and Deacetylation (NuRD) complex is an abundant and highly-conserved complex regulating cell fate transitions and differentiation in many different organisms and developmental contexts *(2–4)*. The multi-protein complex combines two enzymatic activities: lysine deacetylation mediated by Histone Deacetylase (HDAC) 1 and 2 proteins, and ATPase-dependent nucleosome remodeling by Chromodomain Helicase DNA binding protein (CHD) 3 or 4 *(5–7)*. Additional complex partners are the histone chaperone proteins RBBP4 and 7, the zinc-finger proteins GATAD2a or GATAD2b, two MTA proteins (MTA1, MTA2, and/or MTA3) and CDK2AP1 *(8)*. Additionally, the methyl-CpG binding protein family members MBD2 or MBD3 are essential but mutually exclusive NuRD complex members, therefore assembling distinct MBD2-NuRD or MBD3-NuRD complexes *(4, 9)*. Recent structural and biochemical data support the notion that the MBD2 and MBD3 proteins function as a link between the MTA:HDAC:RBBP core and the peripheral GATAD2:CHD:CDK2AP remodeling module *(10–12)*. Absence of MBD2 or MBD3 therefore disrupts NuRD complex functionality. In addition, replacement of MBD2 or MBD3 with PWWP2A results in a distinct complex lacking the remodeling module, also called NuDe complex *(13–15)*. *In vivo*, MBD2 seems dispensable for normal mouse development as MBD2 KO mice display only minor phenotypes but are viable and fertile *(4)*. In contrast, MBD3 is required to exit pluripotency and essential for early mammalian development reflected by lethality of MBD3 KO mouse embryos *(4, 16–18)*.

MBD2 and MBD3 are closely related proteins that share almost 80% homology outside the MBD domain and arose by gene duplication from an ancestral MBD2/3 gene that is present in some metazoans *(4, 9, 19)*. MBD2 and MBD3 contain an MBD and a coiled-coil domain (CC) separated by a disordered protein region, with the latter two being important for protein-protein interaction with the NuRD complex *(20–22)*. Whereas the MBD domain of MBD2 shows high affinity for methylated DNA, the MBD3-MBD domain lacks four conserved amino acids required for the recognition of methyl-CpG. In addition, MBD2 contains a N-terminal glycine-arginine (GR) rich stretch that has been implicated in increasing DNA methylation affinity and interactions with the NuRD complex *(9, 23)*. Differential inclusion of these domains result in various MBD2 and MBD3 isoforms, some with cell type or tissue-specific expression *(16, 24–26)*. Three MBD3 isoforms are present in mouse ESCs: The full-length MBD3a isoform, MBD3b with a truncated MBD domain and MBD3c lacking the MBD domain *(16)*. MBD2 also contains three isoforms: the full-length MBD2a, MBD2b lacking the N-terminal GR repeat and MBD2t lacking the C-terminal CC domain. Based on the presence of either MBD2 or MBD3 in the NuRD complex, MBD2-NuRD and MBD3-NuRD are thought to have distinct functional roles during early development. It is speculated that this is mainly due to their differential binding affinity to methylated DNA by the MBD proteins and recruitment of the NuRD complex to distinct genomic sites. The tissue-specific presence of MBD2- or MBD3-isoforms are expected to further increase the complexity of NuRD complex function. Still, little is known about the direct requirement of the individual MBD2 and MBD3 domains for NuRD complex activity during cellular differentiation. Furthermore, differential and overlapping expression levels of MBD2 and MBD3 isoforms in different cellular contexts convolutes our current understanding about the roles of these different NuRD complexes, requiring further investigation.

Here, we took a systematic approach to dissect the functionality of different NuRD complex compositions during neuronal commitment and terminal differentiation through controlling the expression of MBD2- or MBD3-isoforms. Towards this, we combined neuronal differentiation of engineered murine ES cells with FACS-based measurements of cell identity and transcriptional profiling. In our approach, successful lineage commitment is a direct measurement of a functional NuRD and the role of specific isoforms. While MBD3 is a critical NuRD complex member allowing neuronal differentiation, we show that it functions independent of its MBD domain. Additionally, full-length MBD2 is able to partially compensate MBD3 function. In absence of the GR-stretch or DNA methylation binding affinity, this ability is further elevated to fully compensate absence of MBD3, indicating that these properties prevent a complete redundancy to MBD3. In sum, our results combining functional assays with gene expression analysis of a diverse set of MBD constructs, highlight a partial redundancy of MBD2 and MBD3 during cellular differentiation and point to a more structural than instructive function of the specific MBD family members.

## Results

### Establishment of a functional readout for systematic interrogation of MBD2/MBD3-NuRD complexes

To investigate the distinct roles of MBD2 and MBD3 during lineage commitment, we employed a well-established *in vitro* differentiation system of ESCs towards homogenous populations of neural progenitor cells (NPC) and terminal neurons (TN) *(27)* (Figure 1A). First, we explored published microarray expression data *(28)* of several surface proteins at consecutive differentiation stages (ESC, cell aggregate formation (CA) day4, NPC day 8 and TN day2 and day4, respectively) and identified two neuronal surface proteins, CD24a (CD24) and CD56 (also known as NCAM1), as significantly upregulated at the NPC and TN stage, indicating successful neuronal lineage commitment (Figure 1B). We further established a FACS-based readout on NPCs to quantify the expression of those neuronal surface markers as a measure to score the differentiation potential of ESC. In addition, we assessed successful lineage commitment by the total amount of live cells at progenitor and terminal stages of the differentiation protocol.

**Figure 1:**
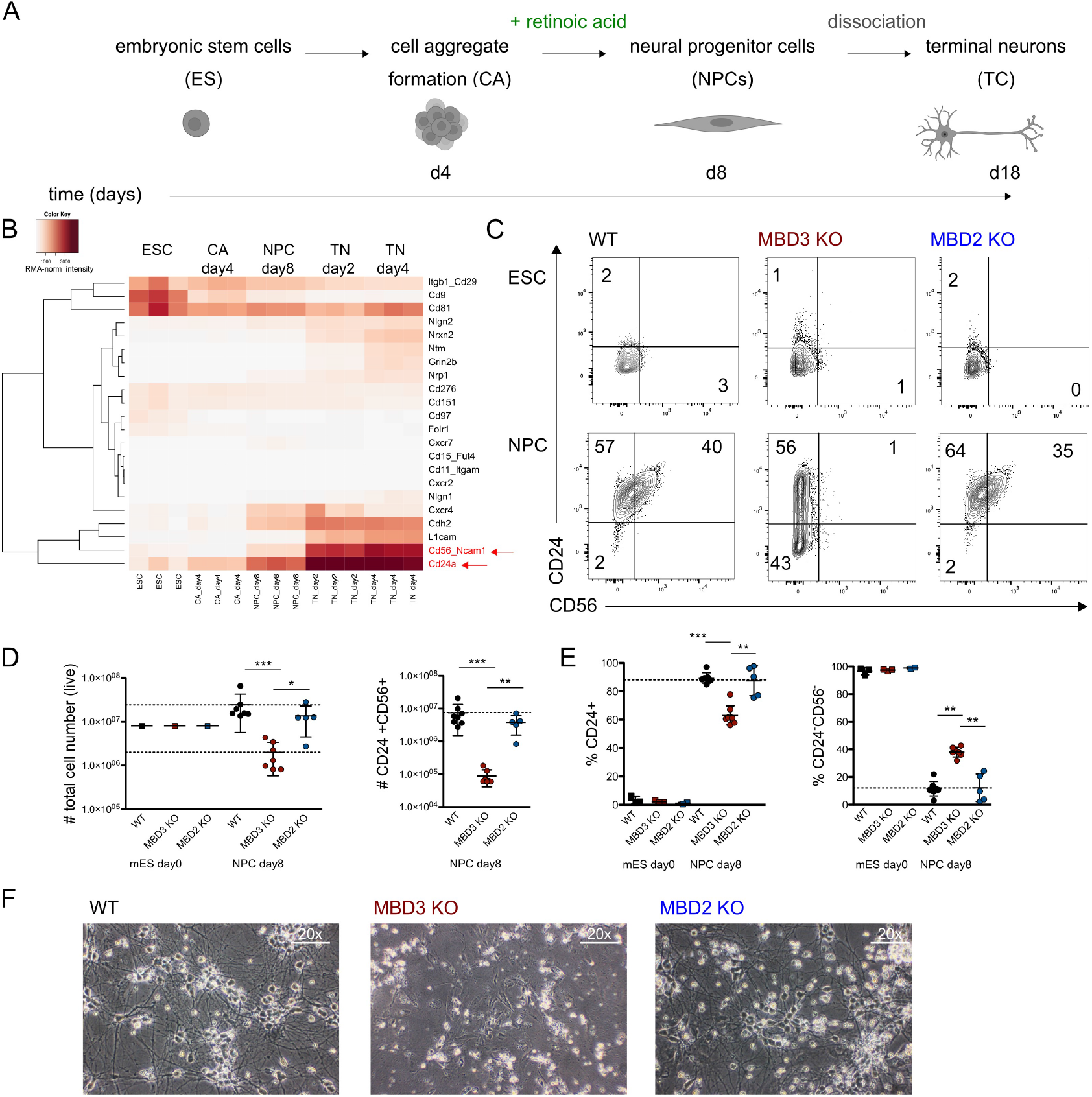
Neuronal differentiation in absence of MBD2 and MBD3. (A) Scheme of *in vitro* ESC differentiation model towards neuronal progenitors (NPCs) and terminal neurons (TN). (B) Heat map showing gene expression of selected surface proteins at different stages of neuronal differentiation. Data is shown in triplicates for ESC, CA day4, NPC day8, TN day2 and TN day4. Shown are RMA-normalized microarray intensity values. **(C)** Representative flow cytometry analysis of CD24 and CD56 surface expression in ES (top) and NPC day8 (bottom) of WT, MBD3 KO and MBD2 KO cell lines. Numbers in quadrants of flow cytometry plots indicate percentages of cells. Flow cytometry analysis showing **(D)** number of live cells (left) and CD24+CD56+ cells (right) and **(E)** percentage of CD24+ (left) and CD24-CD56-(right) in ESC and NPCs day8. Each symbol represents individually-generated cell lines. For each cell line 3 independent clones were analyzed. **P < 0.005 and ***P < 0.0005 (Mann-Whitney two-tailed test). **(F)** Microscopy images of *in vitro* derived neurons from WT, MBD3 KO and MBD2 KO ESCs shown at 20x magnification. Same results were obtained from 2 independent replicates.

We first tested the suitability of this setup on individually-derived MBD2 and MBD3 knock-out (KO) ESC cell lines that were generated in the same genetic background using CRISPR-Cas9. Targeting of MBD2 and MBD3 resulted in a complete loss of MBD2-NuRD or MBD3-NuRD as both MBD2 isoforms (MBD2a and b) and all three MBD3 isoforms (MBD3a, b and c) were not detectable in the respective KO cell lines (Supp. Figure 1A-B). We next differentiated the KO cell lines together with wild type cells and measured CD24 and CD56 levels. Whereas uncommitted ESCs do not express CD24 and CD56 in all three tested lines (WT, MBD2 KO and MBD3 KO), NPCs derived from WT cells expressed either CD24 alone or in combination with CD56 (CD24^+^CD56^+^ double-positive), indicating successful neuronal commitment (Figure 1C). As expected and in line with previous reports *(16)*, MBD3 KO ESCs failed to differentiate towards NPCs, indicated by a more than 10-fold reduction of total number of live cells and CD24^+^CD56^+^ double-positive cells (Figure 1D). Additionally, we detected a significant decrease in the frequency of CD24^+^ cells (89% in WT vs. 61% in MBD3 KO) and a significant increase of uncommitted CD24^−^CD56^−^ cells in MBD3 KO cells, when compared to WT (12% in WT vs. 38% in MBD3 KO) (Figure 1E). Furthermore, MBD3 KO ESCs were not able to form terminal neurons (Figure 1F). Unlike MBD3 KO, MBD2 KO ESCs did not show any noticeable differentiation defects, and similar to the WT ESCs, successfully differentiated towards both NPCs and terminal neurons (Figure 1C-F).

### The MBD domain of MBD3 is dispensable for neuronal differentiation

To systematically test the functional role of different MBD3 isoforms and mutant MBD3 proteins in regulating neuronal lineage commitment, we expressed MBD3 protein variants in MBD3 KO ESCs from a heterologous site and assessed their capacity to rescue the neuronal differentiation phenotype. The MBD3 proteins were expressed from a constitutive promoter, integrated to the same site in the mouse genome via recombinase-mediated cassette exchange (RMCE) *(21)*. This enabled us to generate MBD3 KO ESC lines stably expressing the MBD3 protein variants at comparable levels (Figure 2A). Expression of the MBD3a isoform in MBD3 KO ESC fully rescued the differentiation block to levels observed in WT cells, resulting in comparable amounts of live NPCs, CD24^+^, CD24^+^CD56^+^ double-positive cells and terminal neurons (Figure 2 B-D). The MBD3a isoform represents the full-length MBD3 protein, including the MDB domain. Interestingly, two additional MBD3 isoforms present in ESCs contain a truncated (MBD3b) or completely lack (MBD3c) the N-terminal MBD domain *(16, 24)*. Previous reports suggest that all three isoforms are equally capable of promoting lineage commitment *(16, 29)*. To further evaluate the role of the MBD domain of MBD3 in neuronal differentiation, we generated a MBD3 KO ESC line expressing an MBD3 protein lacking the entire MBD domain (MBD3ΔMBD) (Figure 2A). Similar to the MBD3a full-length variant, MBD3ΔMBD completely rescued the differentiation block of the MBD3 KO ES cells (Figure 2B-D). While there are contradicting reports about the affinity of the MBD3-MBD domain towards methylated-/hydroxy-methylated-CpGs, it is not disputed that the MBD domain of MBD2 binds methylated DNA *(21, 24, 30–32)*. To test if DNA-methylation binding would abrogate MBD3 function during ESC differentiation, i.e. through sequestering MBD3-NuRD to methylated sites, we generated a chimeric MBD3 variant where we replaced the MBD domain with that of MBD2 (MBD3_MBD^MBD2^) (Figure 2A). The MBD3 KO cell line expressing MBD3_MBD^MBD2^ differentiated normally towards the neuronal lineage, resulting in a comparable number of live NPCs and CD24^+^ cells and an equal frequency of CD24^+^ and CD24^+^CD56^+^ cells compared to those observed in WT cultures (Figure 2B-C). Furthermore, terminal neuronal differentiation resulted in an equal number of neurons compared to WT cells (Figure 2D). These results suggest that the MBD domain of MBD3 is not only dispensable for MBD3-NuRD complex function during lineage commitment and neuronal differentiation, but also its replacement with a DNA-methylation sensitive MBD domain does not abrogate MBD3-NuRD function.

**Figure 2:**
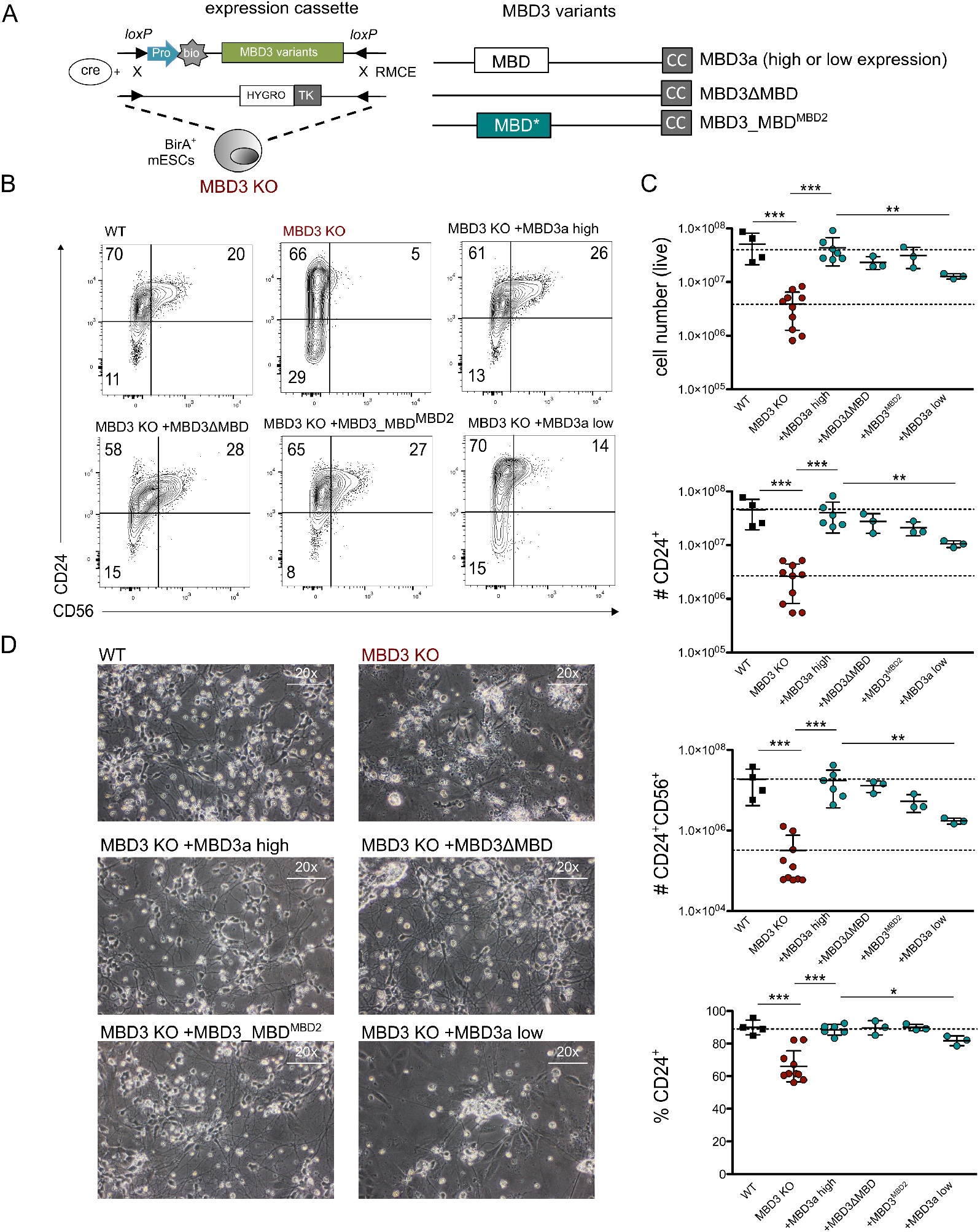
The MBD domain of MBD3 is dispensable for ESC neuronal lineage differentiation. **(A)** Principle of cell line generation via cre recombinase-mediated cassette exchange (RMCE) and following ganciclovir selection. Biotin-tagged MBD3 variant protein cDNAs are targeted to a defined genomic site of the mouse ESC. Introduction of protein variants to the same genomic location enables stable expression and direct comparisons. Triangles: LoxP sites, TK: Thymidne kinase, Pro: promoter either CAG (high expression) or CMV (low expression), bio: biotin acceptor site. **(B)** Representative flow cytometry analysis of CD24 and CD56 surface expression in NPC day8 of WT, MBD3 KO and MBD3 KO stably expressing MBD3a (high or low), MBD3ΔMBD or MBD3_MBDMBD2 variants. Numbers in quadrants of flow cytometry plots indicate percentages of cells. **(C)** Flow cytometry analysis indicating (from top to bottom) the number of live cells, CD24+, or CD24+CD56+ cells, as well as the percentage of CD24+ NPCs at day8 of the differentiation protocol for the cell lines indicated in (B). Each data point represents an individual cell line. For each cell line 3 independent clones were analyzed. **P < 0.005 and ***P < 0.0005 (Mann-Whitney two-tailed test). **(D)** Microscopy images of *in vitro* derived neurons from the cell lines shown in (B) at 20x magnification. Identical results were obtained from 3 independent replicates.

Finally, we tested how MBD3 abundance impacts the differentiation capacity of MBD3 KO ESCs by expressing the full-length MBD3a isoform under a weaker promoter (CMV) (MBD3a^low^) (Figure 2A, Supp. Figure 1C). The lower levels of MBD3a were not sufficient to fully rescue the differentiation block. The MBD3a^low^ cell line resulted in a significant reduction of live NPCs, CD24^+^ and CD24^+^CD56^+^ double-positive cell numbers (4-fold reduction) and a significant reduced percentage of CD24^+^ cells, when compared to WT cultures (90% in WT vs. 78% in MBD3a^low^) (Figure 2B-C). Additionally, MBD3a^low^ cells showed reduced capacity to form terminal neurons (Figure 2D). In sum the results indicated that MBD3 protein abundance rather than its MBD domain composition is crucial for ESC differentiation towards neuronal lineage.

### Full-length MBD2 partially rescues the differentiation block in MBD3 KO ESCs

To understand the role of MBD2-NuRD during lineage commitment we used the same expression system as described above and assessed the capacity of different MBD2 isoforms to rescue neuronal differentiation in MBD3 KO cells (Figure 3A). The generated MBD3 KO ESC lines stably expressed the MBD2 protein variants at comparable levels which allowed us to score their respective function within NuRD (Supp. Figure 2A). To our surprise, heterologous expression of MBD2a could partially rescue the defective lineage commitment of MBD3 KO ESC. We observed a small increase in CD24^+^ and CD24^+^CD56^+^ NPC cell numbers upon MBD2a expression in the MBD3 KO cells (Figure 3B-C). This partial rescue was not observed when introducing the truncated isoform MBD2t that lacks a C-terminal coiled coil (CC) domain required for interactions with the GATAD2:CHD module of the NuRD complex *(20, 21)* (Figure 3A-C), suggesting that the partial rescue observed for the full-length MBD2a requires interactions with the NuRD complex. Despite the low number of CD24^+^CD56^+^ double-positive neuronal progenitors, the MBD2a-expressing MBD3 KO ESCs were able to form terminal neurons (Figure 3D), which is in stark contrast to the absence of neurons in MBD3 KO or MBD2t-expressing MBD3 KO cells (Figures 2D and 3D).

**Figure 3:**
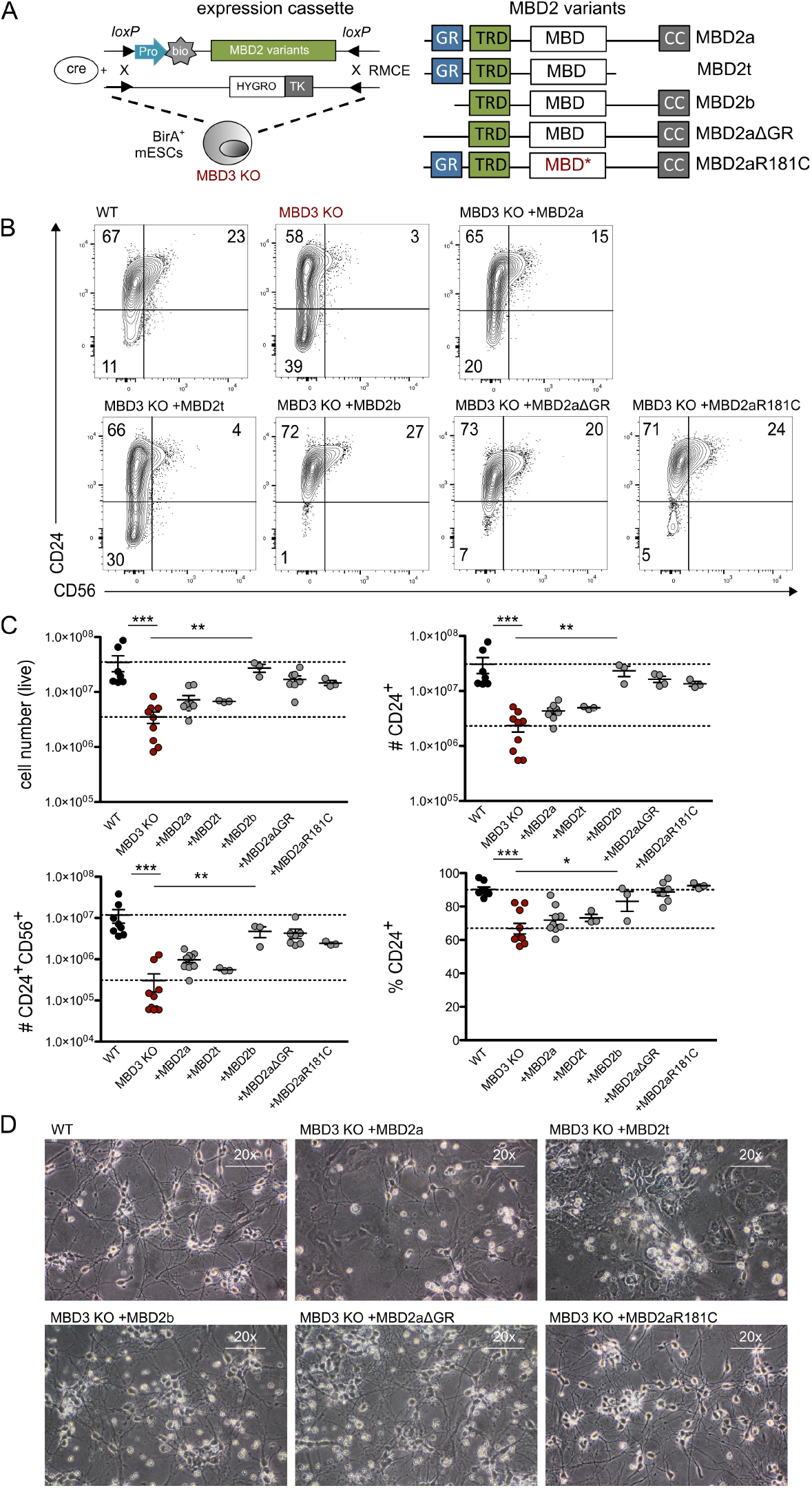
Full-length MBD2 partially rescues the differentiation block in MBD3 KO ESCs. **(A)** Recombinase mediated cassette exchange (RMCE) for MBD2 variants, as in Figure 2A. TRD: transcriptional repressor domain. **(B)** Representative flow cytometry analysis of CD24 and CD56 surface expression in NPCs at day8 of WT, MBD3 KO and MBD3 KO stably expressing MBD2a, MBD2b, MBD2aΔGR, MBD2aR181C or MBD2t. Numbers in quadrants of flow cytometry plots indicate percentages of cells. **(C)** Flow cytometry analysis indicating the number of live cells, CD24+, CD24+CD56+ cells and percentage of CD24+ NPCs at day8 of neuronal differentiation using the cell lines indicated in (A). Each data point represents individual cell lines. For each cell line 3 independent clones were analyzed, except for MBD2t were three technical replicates of one clone were analyzed. **P < 0.005 and ***P < 0.0005 (Mann-Whitney two-tailed test). **(D)** Microscopy images of in vitro derived neurons from cell lines in (A) at 20x magnification. Similar results were obtained from 3-5 independent replicates.

We next wanted to test if other MBD2 isoforms and variants besides MBD2a are able to partially rescue the differentiation block of MBD3 KO ESCs (Figure 3A). First, we introduced the shorter isoform MBD2b that lacks the N-terminal stretch of MBD2a, including the repetitive G/R rich region *(24)*. In contrast to MBD2a, this isoform led to a full rescue of the neuronal differentiation in MBD3 KO ESC at both the NPC and terminal neuron stage (Figure 3B-D). To test a direct contribution of the repetitive G/R stretch, we introduced an engineered MBD2a variant that only lacks the G/R stretch (MBD2aΔGR) (Figure 3A). Similar to MBD2b, heterologous expression of MBD2aΔGR fully restored the neuronal differentiation capacity of MBD3 KO ESC (Figure 3B-D). Finally, we assessed how the mCpG sensitivity of MBD2 contributes to the partial redundancy observed for MBD2a. Towards this, we introduced a known mutation in the MBD domain (R181C) that abrogates its affinity towards methylated DNA *(33)* (MBD2aR181C). Expression of MBD2aR181C in MBD3 KO cells rescued the neuronal differentiation block similar to levels observed in WT cells (Figure 3B-D). Taken together, these results show that MBD2 can partially overcome the differentiation block in MBD3 KO cells, indicating redundant roles for MBD2 and MBD3 in the NuRD complex. This partial rescue is not due to lower expression of MBD2a, since we observe similar levels to MBD3 and other MBD2 variants when expressed from the same heterologous site (Supp. Figure 2A-B). Strikingly, the repetitive G/R stretch and the mCpG binding preference of MBD2 seem to prevent a complete redundancy to MBD3.

### Neuronal gene expression signatures can be restored upon MBD2 or MBD3 reintroduction in MBD3 KO neuronal progenitor cells

Having showed that MBD2 variants are capable of rescuing loss of MBD3 function during neuronal differentiation at different capacities, we next wanted to obtain a better insight into the gene expression programs regulated by the different MBD2/3-NuRD versions introduced in this system. Therefore, we performed RNA-seq at the NPC stage from MBD3 KO lines stably expressing MBD2 and MBD3 variants. Differential gene expression analysis between WT and MBD3 KO cells reveals drastic changes in gene expression with 1821 genes upregulated and 1102 genes downregulated in the MBD3 KO NPCs (Figure 4A). In accordance with the FACS analysis, these changes are completely reverted upon reintroduction of MBD3a, indicating a full rescue of the MBD3 KO phenotype (Figure 4B). We performed similar gene expression analysis for all other cell lines generated in this study. Overall, we observe that cell lines showing a rescue of the MKD3 KO phenotype based on FACS measurements, displayed none or very few deregulated genes compared to wild type NPCs (Figure 4C and Supp. Figures 3A-B). These cell lines included MBD3ΔMBD, MBD2aΔGR, MBD2aR181C. MBD3 KO NPC cells expressing MBD2a had a high degree of deregulated genes (859 up / 1008 down), and MBD2t (2079 up / 1578 down), in line with partial or full failure to rescue differentiation, respectively (Figure 4D and Supp. Figure 3C). Comparing the global gene expression profiles by multidimensional scaling further indicates similarities in gene expression between wild type NPCs and MBD3 KO NPCs expressing MBD3a, MBD3ΔMBD, MBD2aΔGR and MBD2aR181C (Figure 4E). The cluster of these cell lines is separated from MBD3 KO expressing MBD2t that show a full differentiation block, while the MBD3 KO cells expressing MBD2a form an outgroup that is nearer to cell lines failing to fully differentiate (Figure 4E). Clustering of all replicates based on genes differentially expressed between WT and MBD3 KO NPCs further indicates the similarity in gene expression between cell lines that fully rescue the MBD3 KO differentiation block, while cell lines expressing MBD2a showed a correlation to both, MBD3 KO and WT groups (Supp. Figure 3D-E). Focusing on the relevant lineage markers, we observe that cell lines expressing MBD3, MBD3ΔMBD, MBD2aΔGR, MBD2aR181C are able to up-regulate neuronal markers like Neurog1, Neurod4 and Pax6 whereas MBD3 KO lines that express MBD2a or MBD2t maintain a pluripotent signature with high expression of embryonic stem cell-specific genes like Pou5f1 (Oct4), Nanog and Klf4, similar to MBD3 KO cells (Figure 4F).

**Figure 4:**
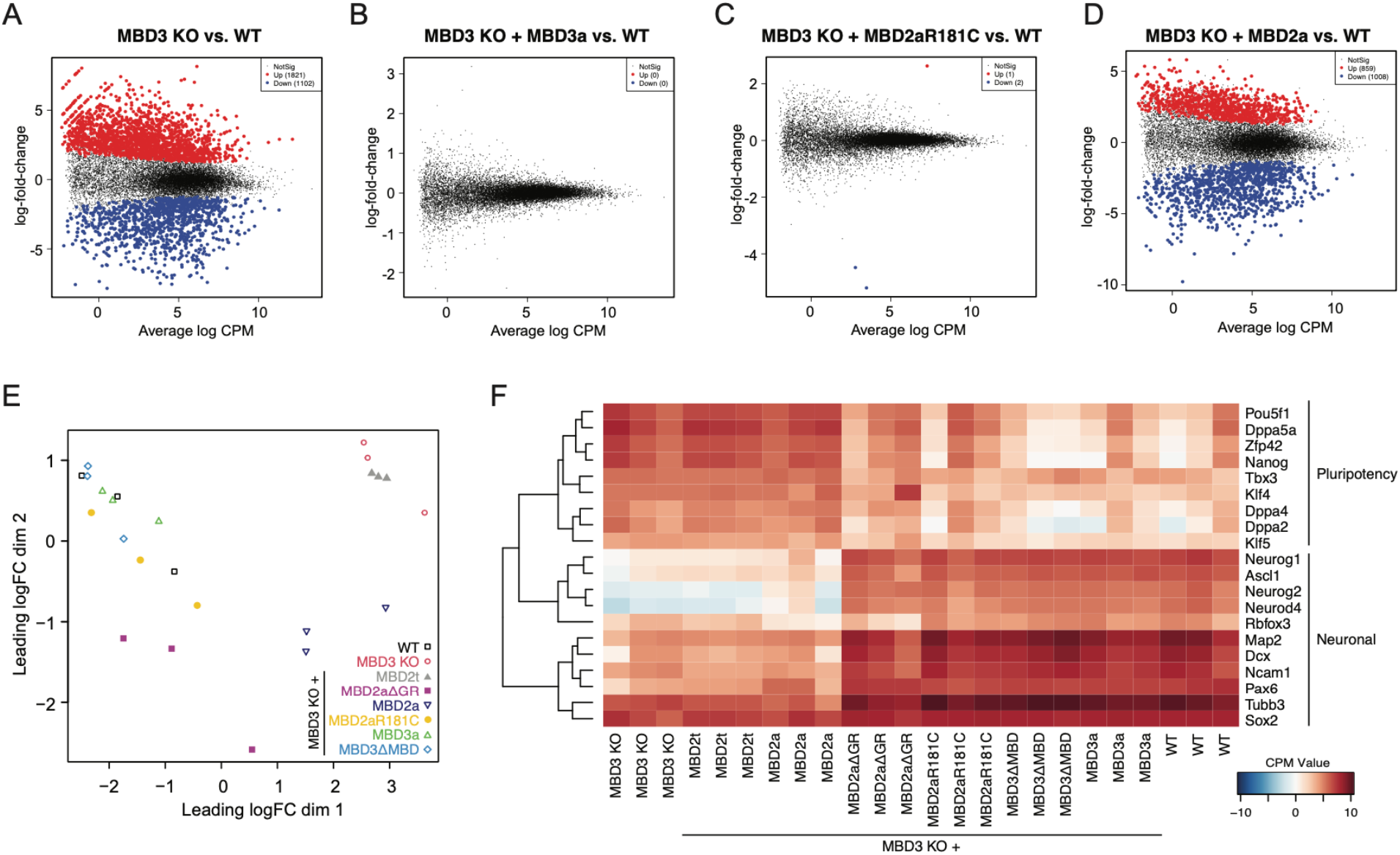
Neuronal gene expression signatures are restored upon expression of mutated MBD2 or MBD3 in MBD3 KO neuronal progenitor cells. **(A-D)** MA-plots showing differential gene expression between MBD3 KO and WT NPCs (A), MBD3 KO + MBD3a vs WT NPCs (B), MBD3 KO + MBD2aR181C vs WT NPCs (C) and MBD3 KO + MBD2a vs WT NPCs (D). Red and blue dots indicate genes with significant changes in gene expression (edgeR, log2FC > 1 | < −1 and adjusted p-value <0.05). **(E)** Multi-dimensional scaling (MDS) plot indicates the degree of similarity for all RNA-seq datasets obtained from NPCs at day8. Each point represents an individually derived clone except for MBD3 KO + MBD2t, were triplicates of one clone were analysed. **(F)** Heatmap indicating gene expression by RNA-seq of pluripotency-associated and neuronal-specific genes in the analyzed samples. Shown in CPM values.

## Discussion

Here we provide a systematic dissection of the different MBD2 and MBD3 isoforms and their protein domains during ESC differentiation. In contrast to other tissues where specific MBD isoforms are present, ESCs express all six MBD2/3 variants (MBD2a,b,t and MBD3a,b,c), which can be mutually exclusively incorporated into the NuRD complex, ultimately forming distinct assemblies with different functionalities *(16, 25)*. NuRD plays an essential role during lineage commitment regulating the exit from pluripotency and enabling proper lineage differentiation *(4, 16, 34, 35)*. Successful ESC lineage commitment therefore serves as a direct measurement of NuRD complex functionality.

By using a well-defined ESC differentiation model towards NPCs and terminal neurons, we showed that while MBD3 is critical for ESC lineage commitment, as previously described *(16)*, this function is independent of its MBD domain and binding to DNA, irrespective of the CpG methylation state. Surprisingly, we found that MBD2a can partially compensate for the loss of MBD3, leading to the generation of fully-differentiated neurons, although at a low frequency. The functional redundancy was critically dependent on its integration into the NuRD complex, as an isoform lacking the C-terminal coiled-coil domain (MBD2t) failed to compensate for absence of MBD3. Conversely to MBD3, the methyl-DNA-binding affinity of the MBD domain in MBD2 prevented full rescue upon expression in MBD3 KO cells, since a point mutation rendering this domain insensitive to methyl-CpG resulted in full differentiation to neuronal cells, with no apparent transcriptional differences to WT cells. This suggests that the affinity of wild type MBD2a for methylated DNA may recruit the NuRD complex away from unmethylated promoters and enhancers to methylated sites in the genome, therefore preventing establishment of correct gene expression patterns. Alternatively, MBD2 interactions with methylated DNA could impede the complete assembly of the NuRD complex, resulting in a limited pool of functional complexes. Based on recent crosslinking mass spectrometry and/or structural data and modelling of the complete NuRD complex, MBD proteins have been shown to bridge the HDAC and CHD modules *(10–12)*. However, if this role, which involves multiple protein-protein interactions with the MBD domain, is compatible with binding to DNA, remains to be fully elucidated. Interestingly, the MBD2b isoform which has similar affinity for methylated DNA, can fully rescue in absence of MBD3. The same is observed when we replace the MBD domain of MBD3 with a methylation sensitive MBD from MBD2, suggesting that the influence of m-CpG readout by the MBD domains on NuRD complex functionality strongly depends on the MBD protein isoform present in the complex. Interestingly, the MBD2b isoform lacks the N-terminal MBD2-specific GR-rich repeat, which is proposed to be methylated by the arginine methyltransferases PRMT1 and PRMT5 and influence mCpG-affinity and incorporation of MBD2 to the NuRD complex *(9, 23)*.

Taken together, the differences observed for the MBD2 isoforms point to a specialized role of these variants in regulating MBD2-NuRD function. Chromatin remodeling complexes often show protein subunit diversity that conveys a specialized function of particular sub-complexes *(1, 36)*. Several studies highlight that NuRD cellular function indeed depends on alternate usage of Mbd2/3, Chd3/4/5 and Mta1/2/3, as MBD2-NuRD but not MBD3-NuRD regulate fetal-hemoglobin switch in adult erythroid cells *(37)* and different CHDs subunits regulate neuronal differentiation and migration with a limited protein redundancy *(38)*. The competition between the MBD2 isoforms with MBD3 proteins for other NuRD components results in different assemblies with – depending on the MBD variant levels present in the analyzed tissue – different functional properties. This can for example lead to the presence of incomplete NuRD complexes lacking the GATAD2:CHD:CDK2AP1 chromatin remodeling module – as in the case of MBD2t or the newly identified component PWWP2A that replaces MBD2/MBD3 from the NuRD complex - also called NuDe complex *(13–15, 20)*. This can also result in differential localization of the NuRD complex to genomic sites based on DNA methylation readout by MBD2. While DNA methylation-dependent localization of MBD2 and localization of MBD3 to unmethylated, active regulatory sites have been reported by multiple groups, other NuRD complex members were predominantly found to localize to the latter, with little overlap to DNA-methylated sites *(21, 34, 35, 39–41)*. It remains to be investigated if different MBD2-isoforms lead to the assembly of alternative NuRD (sub-)complexes with distinct genomic localization or display NuRD-independent functions. Taken together, our data highlight a more complex role of MBD2 isoforms and domains in ESC lineage commitment than previously anticipated.

## Materials and Methods

### Cell culture, cell line generation and neuronal differentiation of embryonic stem cells

Mouse embryonic stem cells (HA36CB1, 129×C57BL/6) were cultured as previously described *(21)*. MBD protein expression constructs in pL1-CAGGS-bio-MCS-polyA-1L or pL1-CMV-bio-MCS-polyA-1L were generated in *(21)*. Subcloning of MBD variants without specific domains were achieved by subcloning from initial plasmids using Gibson-Assembly. MBD protein variant expressing cell lines in MBD3 KO ES cells were obtained by RMCE as previously described *(21)*. Briefly, RMCE constructs were co-transfected with a Cre recombinase expression plasmid (1:0.6 DNA ratio) into RMCE-competent and biotin ligase (BirA)-positive mESCs (HA36CB1)*(21)*. After 10 days ganciclovir selection (3mM), individual clones were picked, and construct integration confirmed by PCR and immunoblotting. The MBD2 KO and MBD3 KO cell lines were generated by co-transfecting pX330-U6-Chimeric_BB-CBh-hSpCas9 (Addgene 42230) with two sgRNA targeting exon 1 for MBD2 KO cell line (sg1_GACTCCGCCATAGAGCAGGG, sg2_CCCCCCCGGATGGAAGAAGG) and exon 2 and exon 3 for MBD3 KO cell line (sg1_CAACTGGCACGTTACCTGGG, sg2_CACCAACCACCCCAGCAACA), respectively. pRR-Puro recombination reporter *(42)* (Addgene 65853) was co-transfected and 36 hours after transfection, cells were treated with 2 μg/ml puromycin for 36 h. Positive KO clones were validated by Sanger sequencing and western blotting. Transfections were conducted using Lipofectamine 3000 reagent (L3000015, Thermo Fisher Scientific) at a 2:1 Lipofectamine/DNA ratio in OptiMEM (31985070, Thermo Fisher Scientific). Neuronal differentiation of embryonic stem cells was performed as previously described *(27)*. Microscopy images were taken at 20x magnification using an Olympus CKX31 microscope and a Canon Eos 550D Camera. Image contrast was increased for better visualization.

### Flow cytometry

For CD24 and CD56 measurements in neuronal progenitors, single-cell suspensions were obtained from neuronal progenitors after 8 days of differentiation, as previously described *(27)*. For cell-surface staining, cells were incubated for 30 min at 4 °C with a saturating concentration of anti-CD24a monoclonal antibody (eBioscience, clone M1/69) and anti-CD56 monoclonal antibody (BD Biociences, clone 809220) in the presence of anti-CD16/CD32 (eBioscience). LIVE/DEAD Fixable Near-IR Dead Cell Stain (L34975, Invitrogen) was used to discriminate cell viability. Samples were acquired using a FACSFortessa (BD Biosciences), and data were analyzed using FlowJo software (Tree Star).

### Immunoblotting

Crude nuclear extracts cells were obtained as described in *(43)*. Membranes were blocked with 5% milk or 5% BSA for detection with antibodies or Streptavidin-HRP, respectively. Primary antibodies against MBD2 (1:2000, ab188474, Abcam), MBD3 (1:2000, ab157464, Abcam), MTA2 (1:2500, sc-9447, Santa Cruz Biotechnology) or anti-LAMIN B1 (1:1000, sc-374015, Santa Cruz Biotechnology) were used overnight at 4°C. Protein detection was facilitated using species-specific antibodies conjugated to horseradish peroxidase and Pierce^®^ Peroxidase IHC Detection Kit (Thermo Scientific) or species-specific secondary antibodies with IRDye Fluorescent Dyes (IRDye 800CW goat anti-rabbit IgG, 1:15,000, LI-COR, P/N 925-32211 and IRDye 680RD goat anti-mouse IgG, 1:15,000, LI-COR, P/N 925-68070) were used and bands were visualized using the Typhoon Biomolecular Imager (Amersham). Biotin detection was performed with Streptavidin-HRP. Molecular weights are indicated by PageRuler Plus Prestained Protein Ladder (Thermo Fisher Scientific).

### Surface marker detection using microarray analysis

Affymetrix Mouse Gene 1.0 ST Array datasets were obtained from GSE27245. Microarrays were RMA-normalized using R/Bioconductor. Probe sets were linked to MGI Gene IDs and probe sets corresponding to surface markers were selected. RMA-normalized values were used to visualize expression of selected surface markers using the heatmap function in R.

### Poly-A RNA-sequencing and differential gene expression analysis

Total RNA was isolated from NPCs using the RNeasy Plus mini kit (Qiagen). RNA integrity was measured using a model 2100 Bioanalyzer (Agilent). PolyA-tailed mRNAs were isolated and enriched using NEB Next Poly(A) mRNA Magnetic Isolation Module according to manufacturer’s instructions. Libraries for 1 μg mRNA were prepared using NEB Next UltraTM II Directional RNA Library Prep Kit for Illumina. Sequencing of library pools and read processing were performed on Illumina NovaSeq according to Illumina standards, with 150-bp single-end sequencing. Sequencing reads were trimmed using trimgalore to remove adapter sequenced and aligned using STAR *(44)* using standard options based on the gene transcript annotation gencode.mouse.v1.annotation.gtf (NCBIM37, mm9). Gene counts were obtained using qCount() in QuasR *(45)* and differential gene expression was performed using the edgeR package with significance set to p-value < 0.05 and log fold change > |1| *(46)*. MA and MDS plots were generated with the plotMD() and plotMDS() functions in edgeR. Heatmap representing gene expression changes for selected genes or all genes differentially expressed between WT and MBD3 KO cells were generated using the gplots::heatmap.2() function using log2-transformed, normalized CPM counts (prior.count = 1).

## Acknowledgements

We thank members of the Baubec laboratory for their input and criticism. Furthermore, we thank members of the Functional Genomics Centre Zurich for their genomics and proteomics support. This work was supported in part by postdoctoral fellowships from EMBO and University of Zurich to N.S., the Baubec laboratory is supported by the Swiss National Science Foundation through SNSF Professorship (183722) and SNSF Sinergia (180354) grants, by the European Research Council (865094 - ChromatinLEGO - ERC-2019-COG), and the EMBO Young Investigator program.

## Author contributions

N.S. and T.B. conceived and designed the study. N.S., I.D. and T.B. developed tools and protocols. N.S. and J.B. generated cell lines and performed experiments. N.S. and T.B. analyzed data. N.S. and T.B. wrote the manuscript with input from all authors.

## Supplementary Figures

**Supplementary Figure 1:**
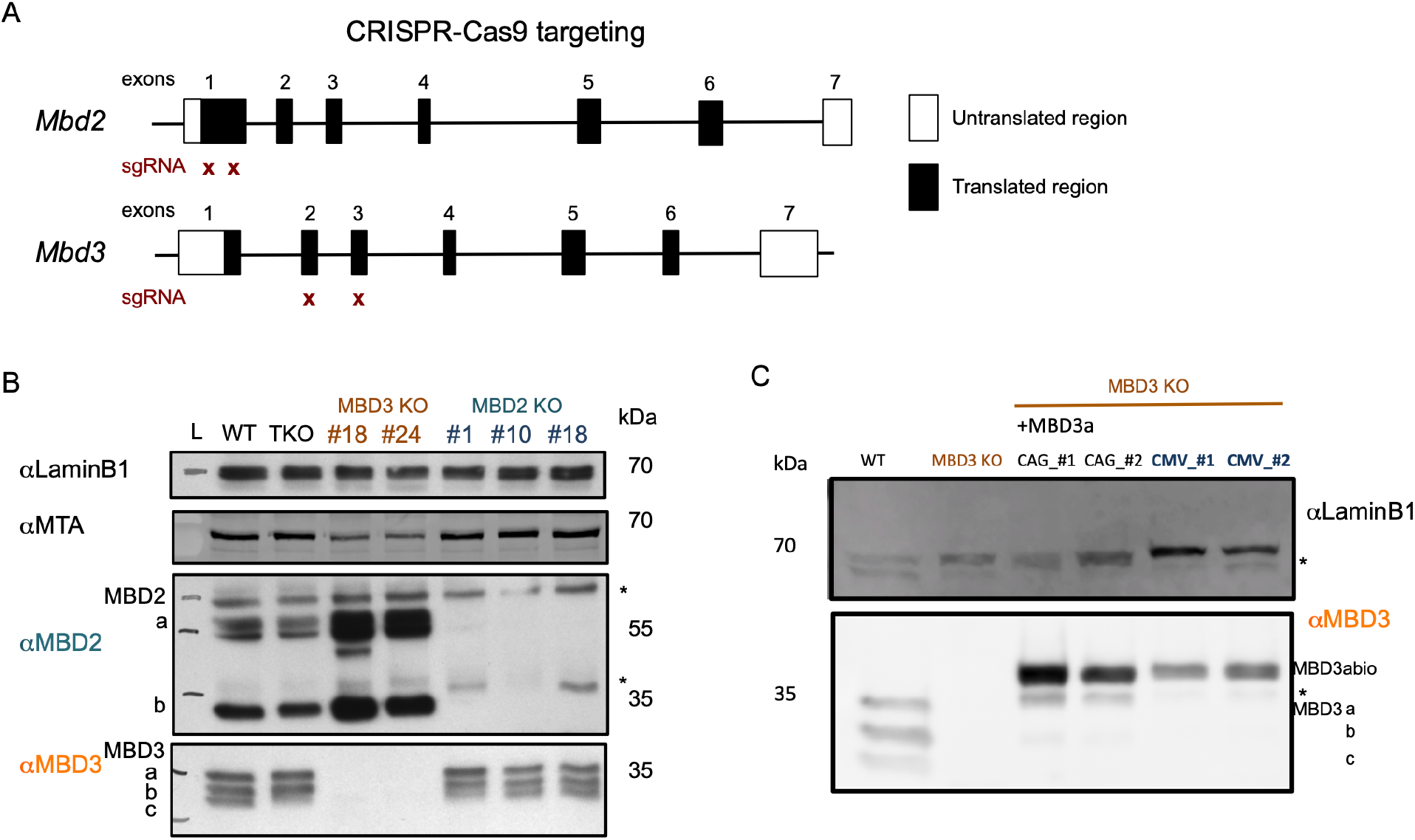
**(A)** Scheme of CRISPR-Cas9 based targeting strategy to generate MBD2 and MBD3 knock-out ESC lines. Both KO cell lines were generated with a two sgRNA targeting approach. Location of sgRNA binding is indicated below. **(B)** Western blot validation using nuclear extracts from WT, Dnmt1/3a/3b-triple KO (TKO) and two MBD3 KO and three MBD2 KO ES cell lines probed with antibodies against MBD2 and MBD3. LaminB1 and MTA act as loading controls. Individual MBD2 and MBD3 isoforms are indicated. Asterisks denote unspecific bands and L denotes the molecular weight marker. **(C)** Western blot indicating MBD3 levels in WT, MBD3 KO, MBD3 KO +MBD3a driven by either CAG or CMV promoters probed with antibodies against MBD3. Two individualy-derived lines are shown for both constructs. LaminB1 acts as a loading control. Approximate sizes are indicated in kDa. Asterisks denote unspecific bands.

**Supplementary Figure 2:**
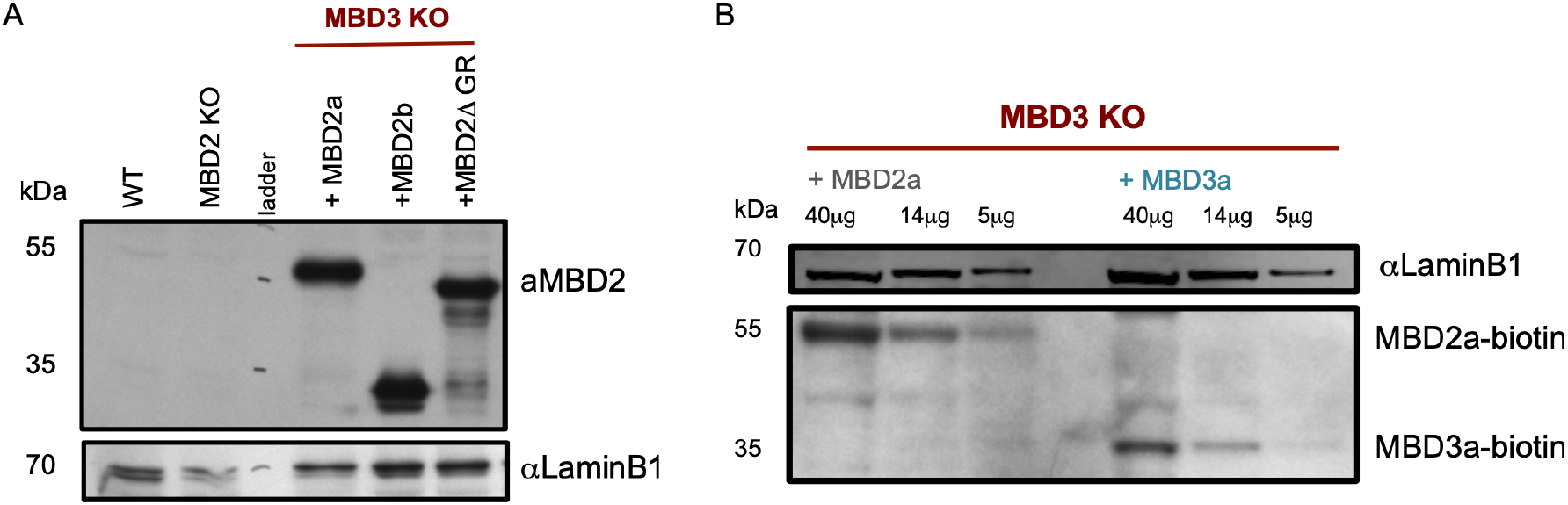
**(A)** Western blot validation of MBD2 KO and MBD3 KO + MBD2aΔGR, +MBD2b, + MBD2a ESC lines probed with antibodies against MBD2. LaminB1 acts as a loading control. Approximate sizes are indicated in kDa. **(B)** Western blot validation of MBD3 KO + biotin-tagged MBD3a or + biotin-tagged MBD2a ESC lines probed with streptavidin-HRP to indicate comparable expression levels of MBD2 and MBD3 from the RMCE site. Various amount of protein extracts (40μg, 14μg, 5μg) were loaded. LaminB1 acts as a loading control. Approximate sizes are indicated in kDa.

**Supplementary Figure 3:**
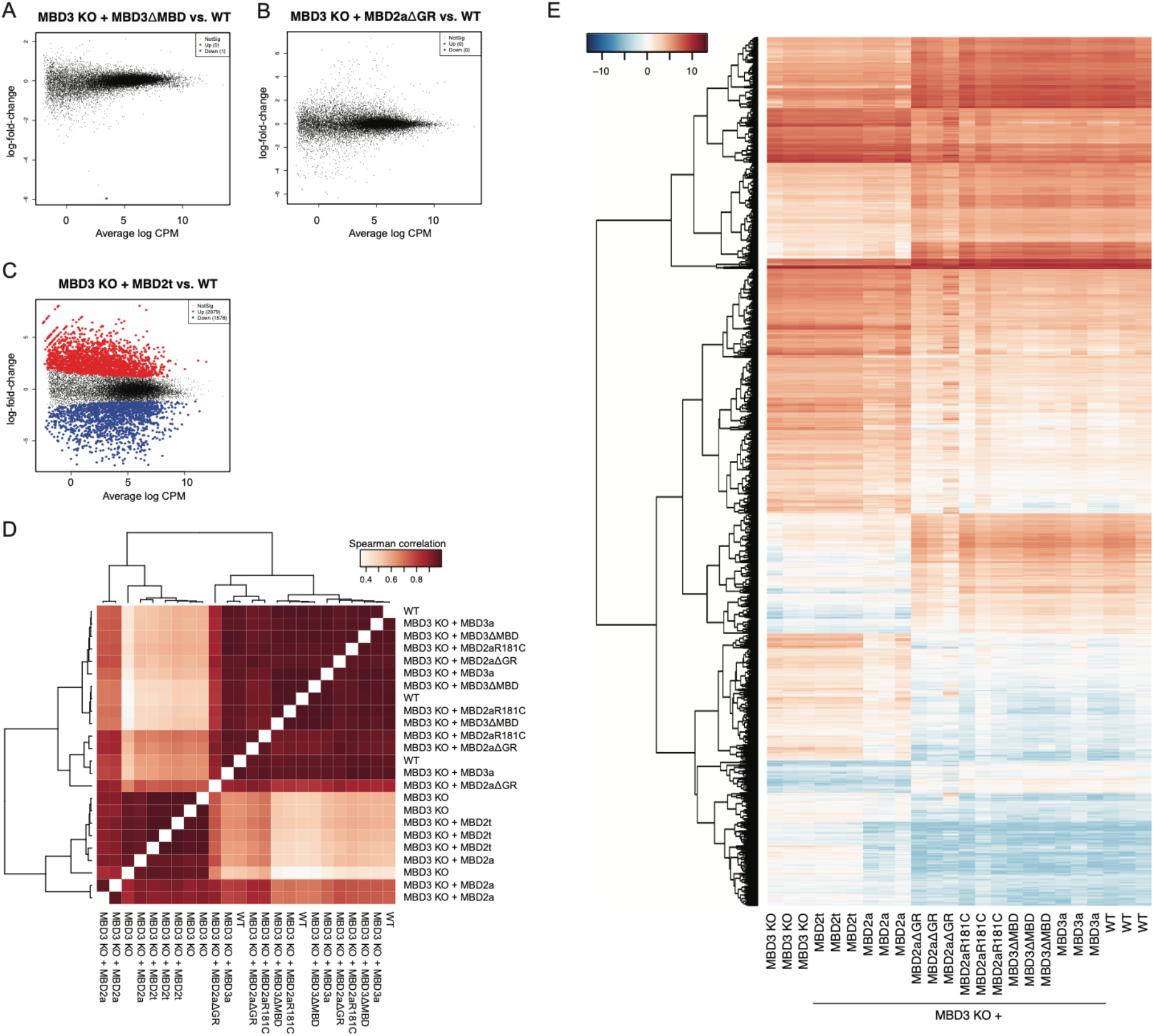
**(A-C)** MA-plots showing differential gene expression between MBD3 KO + MBD3ΔMBD and WT NPCs (A), MBD3 KO + MBD2aΔGR vs WT NPCs (B), and MBD3 KO + MBD2t vs WT NPCs (C). Red and blue dots indicate genes with significant changes in gene expression (edgeR, log2FC > 1 | < −1 and adjusted p-value <0.05). **(D)** Cross-correlation matrix and unsupervised clustering based on genes differentially expressed between WT and MBD3 KO NPCs (Figure 4A), indicates similarities in gene expression between the analyzed cell types. Spearman rank correlation is shown. **(E)** Unsupervised clustering of all differentially expressed genes between WT and MBD3 KO NPCs shows their expression in the analyzed samples. Three individually-derived clones were analyzed for all indicated cell lines except for MBD3 KO +MBD2t were triplicates of one clone where analysed.

